# Did a novel virus contribute to late blight epidemics?

**DOI:** 10.1101/383653

**Authors:** Guohong Cai, Kevin Myers, William E. Fry, Bradley I. Hillman

## Abstract

*Phytophthora infestans* is the causal agent of potato and tomato late blight. In this study, we characterized a novel RNA virus, Phytophthora infestans RNA virus 2 (PiRV-2). The PiRV-2 genome is 11,170 nt and lacks a polyA tail. It contains a single large open reading frame (ORF) with short 5’- and 3’-untranslated regions. The ORF is predicted to encode a polyprotein of 3710 aa (calculated molecular weight 410.94 kDa). This virus lacks significant similarity to any other known viruses, even in the conserved RNA-dependent RNA polymerase region. Comparing isogenic strains with or without the virus demonstrated that the virus stimulated sporangia production in *P. infestans* and appeared to enhance its virulence. Transcriptome analysis revealed that it achieved sporulation stimulation likely through down-regulation of ammonium and amino acid intake in *P. infestans*. This virus was faithfully transmitted through asexual reproduction. Survey of PiRV-2 presence in a *P. infestans* collection found it in most strains in the US-8 lineage, a very successful clonal lineage of *P. infestans* in North America. We suggest that PiRV-2 may have contributed to its success, raising the intriguing possibility that a potentially hypervirulent virus may contribute to late blight epidemics.

**Author Summary:** Potato late blight, the notorious plant disease behind the Irish Potato Famine, continues to pose a serious threat to potato and tomato production worldwide. While most studies on late blight epidemics focuses on pathogen virulence, host resistance, environmental factors and fungicide resistance, we present evidence in this study that a virus infecting the causal agent, *Phytophthora infestans*, may have played a role. We characterized a novel RNA virus, Phytophthora infestans RNA virus 2 (PiRV-2) and examined its effects on its host. By comparing identical *P. infestans* strains except with or without the virus, we found that PiRV-2 stimulated sporulation of *P. infestans* (a critical factor in late blight epidemics) and increased its virulence. We also profiled gene expression in these strains and identified potential molecular mechanisms through which PiRV-2 asserted its sporulation stimulation effect. In a survey of PiRV-2 presence in a *P. infestans* collection, we found PiRV-2 in most isolates of the US-8 clonal lineage, a very successfull ineage that dominated potato fields in North America for several decades. We suggest that PiRV-2 may have contributed to its success. Our findings raise the intriguing possibility that a potentially hypervirulent virus may contribute to late blight epidemics.

## Introduction

Potato late blight caused devastation in the 1840s and led to food shortage across Europe. In Ireland, where the poor were overwhelmingly dependent on potato, it gave rise to the Irish potato famine - over one million people died and many more were forced to flee [1]. Late blight has proved to be a destructive disease that is difficult to manage and control. Even after extensive research efforts for more than one and a half centuries, it continues to devastate potatoes and tomatoes worldwide and is estimated to cause over 6.7 billion dollars annually in yield losses and control costs [2-4]. Late blight can act very rapidly and leave very little time for growers to suppress it once an epidemic has started.

The causal pathogen of potato late blight is named *Phytophthora infestans* (Mont.) de Bary. It also causes late blight of tomato. *P. infestans* is a member of oomycetes, which are morphologically similar to filamentous fungi and occupy similar environmental habitats. Historically, *P. infestans* and other oomycetes were classified as fungi, but they have distinct attributes that separate them from true fungi and in fact are not closely related to fungi. For example, oomycetes have diploid vegetative cells and cellulosic cell walls, while fungi are usually haploid in vegetative state and their cell walls contain chitin. Phylogenetic studies place oomycetes in the major group Stramenopila [5-7], which also includes brown algae and diatoms.

By the mid-twentieth century, late blight was managed to tolerable level through various tools, including cultural practices, host resistance, and fungicide usage. However, by the end of twentieth century, late blight came back with vengeance (reviewed in [8]). The resurgence started in Europe in early 1980s, later in Middle East and Far East, and in North America by late 1980s and early 1990s. The resurgence was caused by multiple independent introductions of exotic and often fungicide-resistant strains from Mexico [9], the center of genetic diversity and origin of *P. infestans*.

Prior to the late 20^th^ century, global populations of *P. infestans* outside Mexico were dominated by a single clonal lineage, US-1 (mating type A1) [10]. It was soon replaced world-wide by new strains [11]. In North America, US-8 was arguably the most successful among the introduced lineages. It is extremely virulent on potato, but not as much on tomato [12]. After first being detected in a single county in New York in 1992, it spread to 23 states during 1994 and 1995, and was found in most potato production regions in the United States and part of Canada by 1996 [13]. Many other clonal lineages, up to US-24, have been reported before and after US-8 [14, 15]. Most lineages have come and gone, but US-8 has proved to be enduring.

Research on late blight epidemics usually focuses on pathogen virulence, host resistance, environmental factors and fungicide resistance, but any potential role of virus infection of *P. infestans* has not been examined. We previously surveyed *P. infestans* isolates and reported the discovery of four RNA viruses in this organism [16]. Three of these viruses have been characterized and their descriptions reported [16-18]. Several additional viruses have been reported in other oomycetes [19]. Like many fungal viruses that are asymptomatic in their hosts, these oomycete viruses caused no measurable phenotype in their host organisms.

In this paper, we present the characterization of Phytophthora infestans RNA virus 2 (PiRV-2), a novel RNA virus with a genome of 11,170 nt and no close affinity to other known viruses. This virus stimulated sporangia production and appeared to enhance virulence of *P. infestans*. Transcriptome analysis revealed that PiRV-2 likely stimulated sporulation through restriction of ammonium and amino acid intake. Survey of PiRV-2 presence in *P. infestans* population suggested that this virus may have contributed to the success of the US-8 lineage and played a role in late blight epidemics.

## Results

### Nucleotide sequence analysis of PiRV-2

Two *P. infestans* isolates in the US-8 lineage, US940480 and US040009, were previously found to harbor PiRV-2 dsRNA [16]. Repeated effort failed to yield virus-like particles (VLPs) (not shown). DsRNA from US940480 was sequenced by random RT-PCR, RT-PCR with sequence-specific primers, and RNA ligase-mediated RACE (Supplementary Figure S1). After assembly of the overlapping clones, the viral genome was found to be 11,170 nt, matching previous estimate [16].

Examination of the coding potential of all six reading frames revealed a long open reading frame (ORF) on one strand, designated as the positive strand (Fig. 1). This ORF spanned from nt 7 to 11,139, and could encode a polyprotein of 3710 aa (calculated molecular weight 410.94 kDa). The first AUG codon in the ORF at nt 7–9 had an A at both −3 and +4 positions, predicted to be a more favorable context than the two AUG codons immediately downstream in this reading frame (nt 253–255 and 292–294, respectively)[20, 21], both of which had a pyrimidine (C) at −3 position. Pending experimental evidence, we assumed for sequence analysis purposes the AUG codon at nt 7-9 was the translation initiation site. This implied a short 5’ untranslated region (UTR) of 6 nt and a 3’ UTR of 31 nt. A signal peptide was not identified at the N-terminal region of this protein, but 14 potential transmembrane regions in two clusters were predicted – one in the middle section and one close to the C-terminus (Fig 1). No other significant ORFs were found on either strand of PiRV-2/US940480.

**Fig 1.**
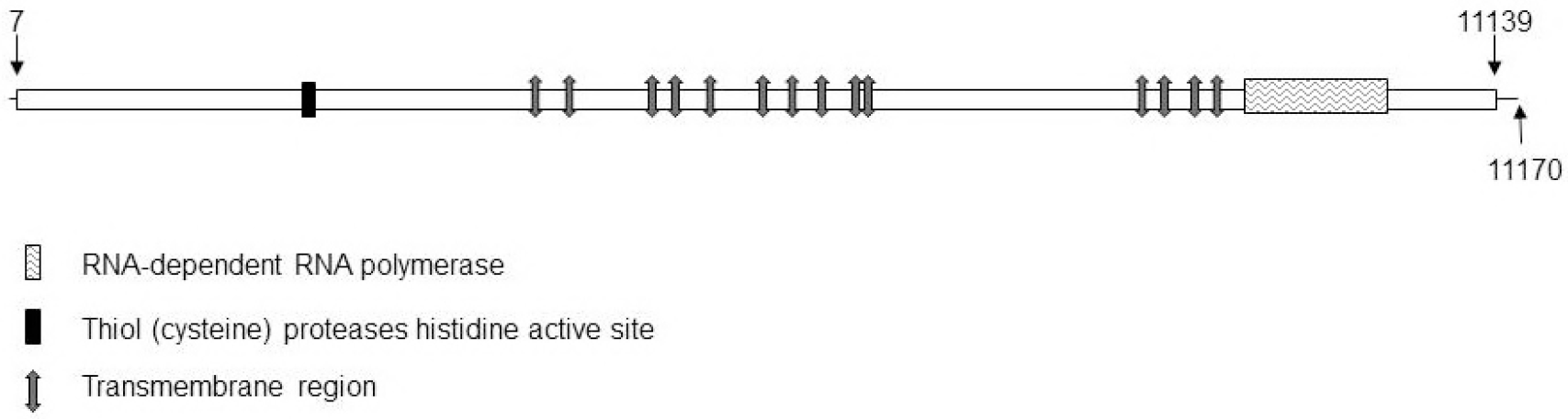
Genome structure of PiRV-2. Lines represent the 5’ and 3’ untranslated regions, the open box represents the predicted single ORF of 11,130 nt beginning at nt 7 and terminating at nt 11,139, and the numbers indicate nucleotide positions. The few predicted functional domains identified on the 3,710 aa protein sequence are provided.

PiRV-2 dsRNA from *P. infestans* isolate US040009 was also sequenced (Supplementary Figure S1). PiRV-2/US940480 and PiRV-2/US040009 shared 99.5% nucleotide identity, with 58 single nucleotide polymorphisms (SNPs) between the two sequences; there were no indels. There were 25 synonymous SNPs while the other 33 were non-synonymous SNPs, and no SNP induced premature termination of the ORF. All downstream analysis was based on PiRV-2/US940480.

### PiRV-2 dsRNA encodes a putative viral RNA-dependent RNA polymerase but lacks similarity to any known viruses

At the nucleic acid level, no significant similarity was found between PiRV-2 and any other sequences in GenBank non-redundant database. When the deduced protein sequence was used as query in BLASTP searches, only marginal similarities were identified between part of PiRV-2-encoded protein and the RNA-dependent RNA polymerase (RdRp) regions of several astroviruses (e.g. *Mamastrovirus 3*, GenBank accession ALA16028, bit score 50.1, E-value 0.15). RdRps of astroviruses contain RdRp_1 domain (pfam00680). Searches against various protein domain profile databases identified an RdRp domain close to the C-terminus of the PiRV-2-encoded protein. For example, in a search of the Conserved Domain Database v3.16 [22], an RdRp (cd01699) was identified at aa 3264–3407 (E-value 3.13e-03). Similarly, scanning Prosite (release 20.120), a database of protein domains, families and functional sites as well as associated patterns and profiles [23, 24], identified PS50507 (RdRp of positive ssRNA virus catalytic domain profile) at aa 3262 – 3392 (score10.724). In contrast, a search of the Pfam database did not find significant similarity between PiRV-2 and any RdRps or other protein domains/families.

All RdRps and many DNA-directed polymerases employ a right-hand shape fold with three subdomains termed fingers, palm and thumb [25]. Only the palm subdomain is well conserved among all of these enzymes. In RdRps, the palm subdomain comprises three well conserved motifs, D-x(4,5)-D, [S, T]-G-x3-T-x3-N and G-D-D [25, 26], where entries in the bracket represent amino acid options in that position, x stands for any amino acid, and numbers represent the number of preceding amino acids. An exception is the virus family *Birnaviridae*, which lacks the G-D-D motif [27]. All three motifs were found in PiRV-2 (Fig 2A). Based on the above data, we concluded that PiRV-2 encoded an RdRp. An alignment of core RdRp regions from representative viruses with RdRP_1 or RdRP_4 domain was built (Fig 2A), and a neighbor joining tree based on this alignment showed that PiRV-2 did not fall into any known virus families (Fig 2B).

**Fig 2.**
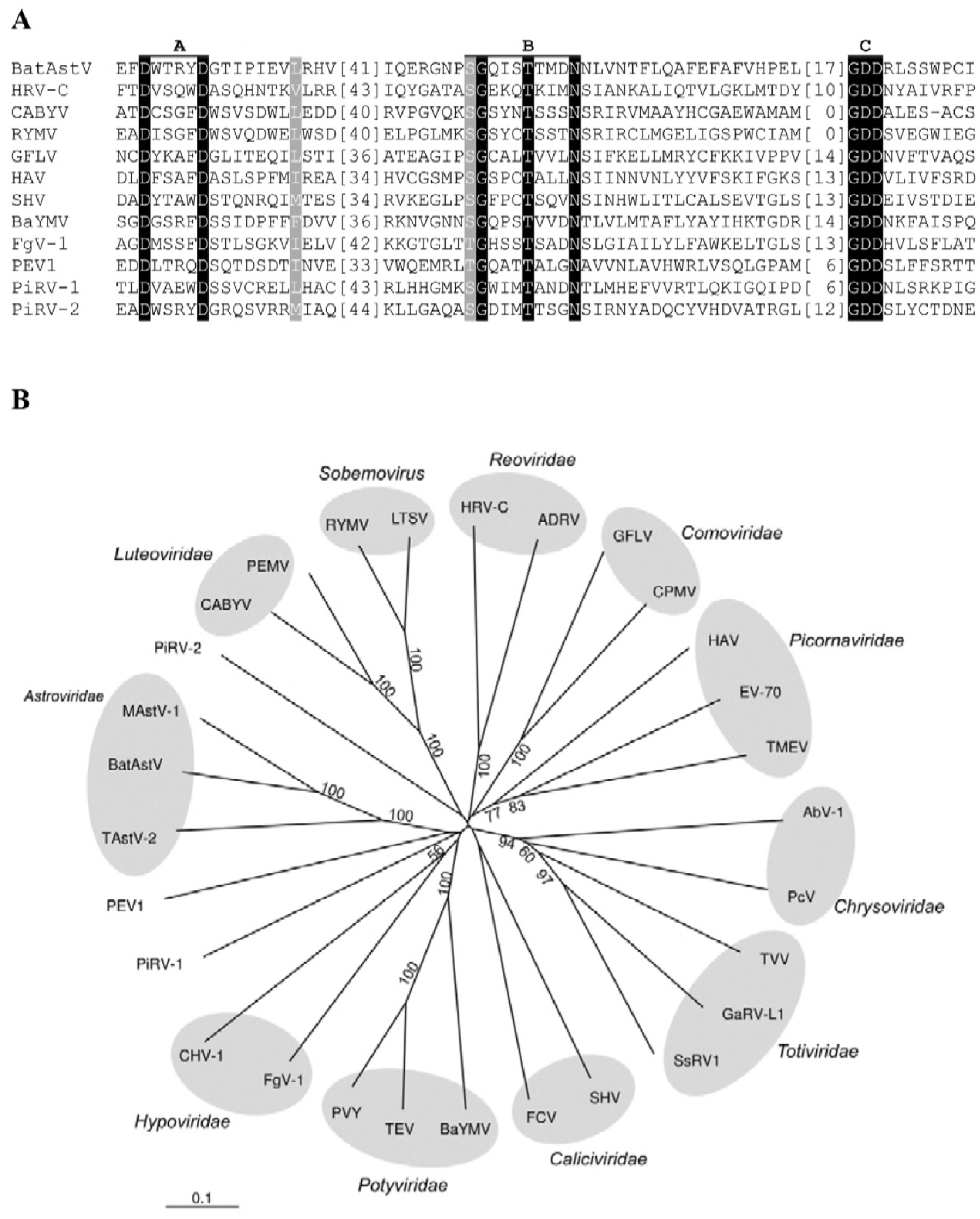
PiRV-2 encodes a predicted viral RNA-dependent RNA polymerase with no close similarity to known viruses. **A.** A subset of the alignment of RdRp regions of PiRV-2 and other RNA viruses, including ssRNA and dsRNA viruses. Three conserved motifs of RdRp palm subdomain, A-C, are labeled on the top. Amino acid residues identical in all viruses are highlighted with a black background and white lettering; those belonging to the same group are highlighted with a gray background and white lettering. The grouping of amino acids based on similarity is as follows: A, G; L, I, V, M, F, Y, W; S, T; K, R; D, E, N, Q. Numbers within the brackets indicate the number of amino acids not shown. **B.** Neighbor-joining tree based on the alignment of RdRps between motifs A and C. Representatives from diverse virus groups were included. A subset of the alignment is shown in **A.** Bootstrap values (1000 replicates) in percentage are labeled for branches with more than 50% support. Full virus names and EMBL/GenBank/DDBJ accession numbers for the protein sequences (in parentheses) are: AbV-1, *Agaricus bisporus virus 1* (Q90193); ADRV, *Adult diarrhea rotavirus* (P35942); BatAstV, *Bat astrovirus* (ACF75855); BaYMV, *Barley yellow mosaic virus* (Q04574); CABYV, *Cucurbit aphid-borne yellows virus* (Q65969); CHV-1, *Cryphonectria hypovirus 1* (AAA67458); CPMV, *Cowpea mosaic virus* (P03600); EV-70, *Human enterovirus 70* (P32537); FCV, *Feline calicivirus* (P27407); FgV-1, *Fusarium graminearum virus 1* (AAT07067); GaRV-L1, *Gremmeniella abietina RNA virus L1* (Q99AT3); GFLV, *Grapevine fanleaf virus* (P29149); HAV, *Hepatitis A virus* (P26580); HRV-C, *Human rotavirus C* (Q91E95); LTSV, Lucerne transient streak virus (Q83093); MAstV-1, *Mink astrovirus 1* (AA032082); PcV, *Penicillium chrysogenum virus* (Q8JVC2); PEMV, *Pea enation mosaic virus* (P29154); PEV1, *Phytophthora endornavirus 1* (CAI47561); PiRV-1, Phytophthora infestans RNA virus 1 (FJ373316); PVY, *Potato virus Y* (Q02963); RYMV, *Rice yellow mottle virus* (Q86519); ScV-L-A, *Saccharomyces cerevisiae virus L-A* (Q87025); ScV-L-BC, *Saccharomyces cerevisiae virus L-BC* (Q87027); SHV, *Southampton virus* (Q04544); SsRV1, *Sphaeropsis sapinea RNA virus 1* (Q9YXE6); TAstV-2, *Turkey astrovirus 2* (AAF60952); TEV, *Tobacco etch virus* (P04517); TMEV, *Theiler’s murine encephalomyelitis virus* (P08544); and TVV, *Trichomonas vaginalis virus* (Q90155).

In addition to the RdRp region, scanning the Prosite database also found the motif signature of eukaryotic thiol (cysteine) proteases histidine active site (PS00639) at aa 744–754 (SYHSVLMAGCS). Eukaryotic thiol proteases have an essential catalytic triad of cysteine, histidine and asparagine. The motif signatures of the cysteine and asparagine active sites were not found, either by scanning Prosite database or by visual inspection. It remains to be determined whether PiRV-2 encodes a protease around the identified histidine active site.

### PiRV-2 stimulates sporangia production in *P. infestans* and enhances its virulence

To examine the effect of PiRV-2 on its host, the virus was cured from isolate US940480 and then re-introduced into the cured isolates through anastomosis. To cure PiRV-2, US940480 was grown on Ribavirin-containing rye agar and transferred by hyphae-tipping. After four iterations, the virus was found to be cured in two cultures, as confirmed by dsRNA extraction and RT-PCR (Supplementary Figure S2). These two cultures were named US940480/PiRV-2C1 and US940480/PiRV-2C2, respectively. To introduce PiRV-2 back into cured isolates, US940480 was co-cultured in close proximity with US940480/PiRV-2C1 or US940480/PiRV-2C2 on rye agar and an agar plug was taken from the side of cured isolates. Virus transmission was confirmed by RT-PCR (Supplementary Figure S3). The resultant cultures were named US940480/PiRV-2T1 and US940480/PiRV-2T2, respectively.

US940480, US940480/PiRV-2C1, US940480/PiRV-2C2, US940480/PiRV-2T1 and US940480/PiRV-2T2 were grown on rye agar to compare their growth rate, colony morphology and sporulation. Slightly but significantly (P =0.05) faster growth rate was observed in the virus-cured isolates. The virus-cured isolates also produced denser aerial mycelium compared to US940480 based on visual observation (Fig. 3A). The most profound difference was observed in sporangia production: virus-infected isolate US940480 produced abundant sporangia, while cured isolates US940480/PiRV-2C1 and US940480/PiRV-2C2 sporulated sparsely (Fig. 3A, inset). On average, the PiRV-2-containing isolate US940480 produced 9 to 125 times as many sporangia as its isogenic, virus-free counterparts US940480/PiRV-2C1 and US940480/PiRV-2C2 did, and the difference was always statistically significant (P = 0.05, Fig 3C and not shown).

**Fig 3.**
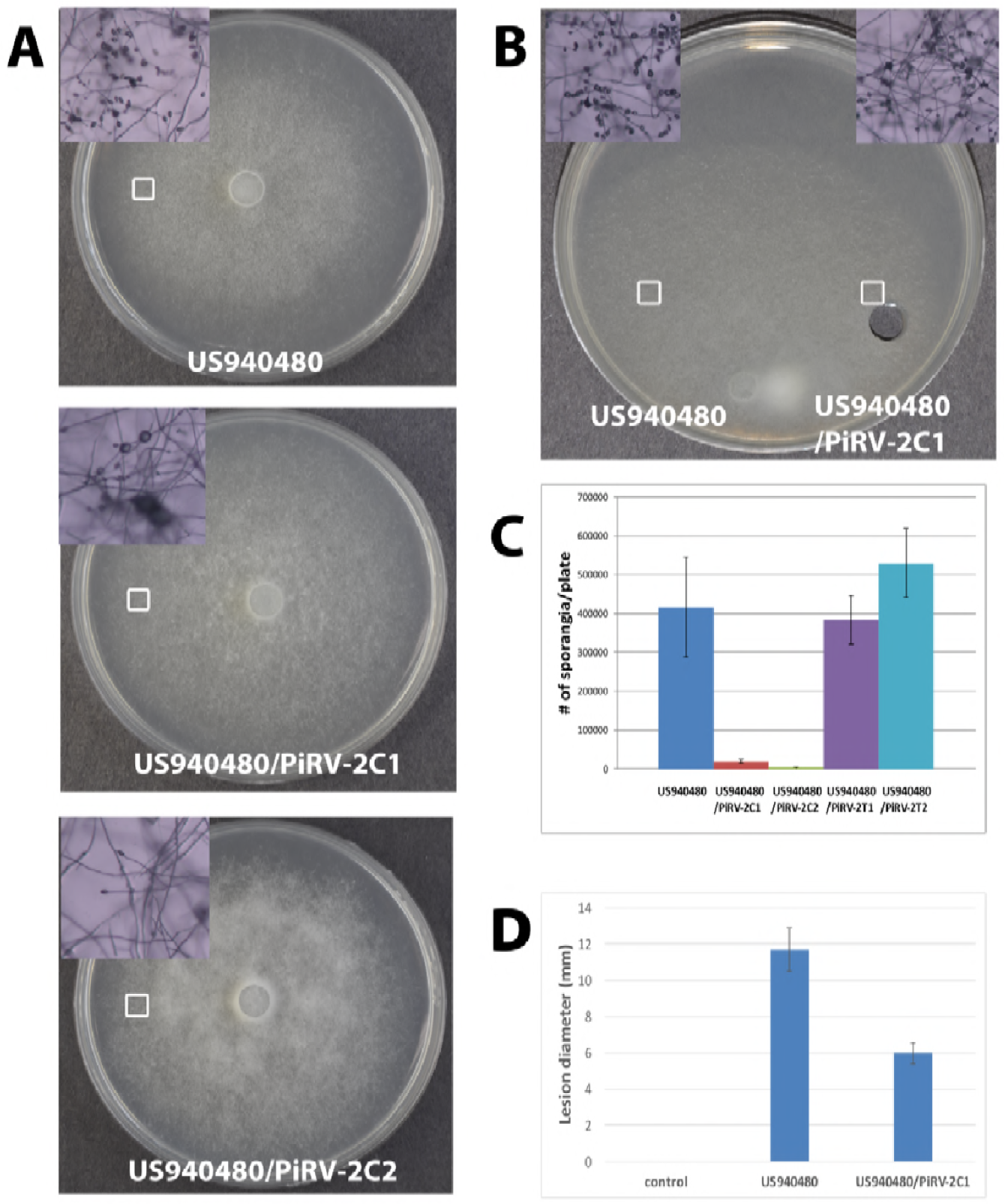
PiRV-2 stimulates sporangia production in *P. infestans* and enhances its virulence. **(A)** Colony morphology and sporulation (insets) of US940480, US940480/PiRV-2C1 and US940480/PiRV-2C2. **(B)** Co-culturing of US940480 and US940480/PiRV-2C1 transmitted the virus into the latter and restored its sporangia production. The white squares in **A** and **B** indicate the areas microscopically observed and photographed. An agar plug was taken from the side of US940480/PiRV-2C1 to confirm virus transmission (supplementary Figure S3). **(C)** Sporangia production in isogenic PiRV-2-containing and PiRV-2-cured isolates. **(D)** Lesion size caused by isogenic PiRV-2-containing and PiRV-2-cured isolates in detached leaf assay. Vertical bars, mean; and vertical lines, standard error. Representative result from one of two experiments.

Co-culturing of US940480 with US940480/PiRV-2C1 or US940480/PiRV-2C2 re introduced the virus into the virus free isolates (Supplementary Figure S3 and not shown) and restored their sporangia production (Fig. 3B-C). Sporangia production in US940480/PiRV-2T1 and US940480/PiRV-2T2 were not significantly different from sporangia production in US940480 (Fig. 3C).

To determine the impact of PiRV-2 on host virulence, US940480 and US940480/PiRV-2C1 were used to infect leaves of susceptible potato cultivar Red La Soda in detached leave assay. Lesions caused by US940480 were significantly larger than those caused by US940480/PiRV-2C1 (Fig. 3D). We were not able to obtain sufficient numbers of sporangia from US940480/PiRV-2C2 to compare those from US940480.

### PiRV-2 causes transcriptome changes in *P. infestans*

To understand the molecular mechanism(s) through which PiRV-2 acted on its host, transcriptomes of US940480 and US940480/PiRV-2C1 were sequenced. Three independent mRNA libraries each for US940480 and US940480/PiRV-2C1 were sequenced on Illumina Miseq platform, generating 5.08 million to 7.04 million reads for each library. The reads were first mapped to the PiRV-2 genome. Two out of three US940480 mRNA libraries contained one and two reads, respectively, that were aligned to PiRV-2. No read from the three US940480/PiRV-2C1 mRNA libraries was aligned to the virus genome. This was in agreement with the fact that US940480 harbored PiRV-2 while US940480/PiRV-2C1 did not. Since PiRV-2 did not have a polyA tail, and mRNA was extracted from total RNA using polyT beads, only trace amount of PiRV-2 RNA was introduced into the libraries.

The sequence reads were then mapped to the genome of *P. infestans* [28] to obtain read counts for individual genes. With a Benjamini-Hochberg adjusted *P* value of 0.01 and a minimum of log_2_ fold change (LFC) of 1 (2-fold change) requirements, 848 differentially expressed genes (DEGs) were identified, with 431 genes up-regulated by PiRV-2 and 417 genes down-regulated by this virus (Supplementary Table S3).

Four groups of genes were enriched in up-regulated DEGs. The first group (i) was histone protein genes. Of the 18 histone protein genes annotated in *P. infestans* genome, 9 were up-regulated, a 20.6-fold enrichment. Two additional histone genes were induced to a lesser extent (padj < 0.05, LFC > 0.5) and none was significantly down-regulated (Supplementary Figure S4A and Supplementary Table S4). Also enriched were (ii) genes for flagellum-related proteins, (iii) proteins with epidermal growth factor (EGF)-like conserved site, and (iv) genes in the glycolytic process (Supplementary Figure S4 B, C and D), with 10.0-, 9.9- and 7.4-fold enrichment, respectively.

Genes in the ribosome pathway were highly represented in the down-regulated DEGs. Of the 110 genes in *P. infestans* that mapped to the KEGG ribosome pathway, 23 were down-regulated, a 9-fold enrichment. An additional 34 genes in this pathway were down-regulated to a lesser extent (padj < 0.05, LFC < −0.5) and no gene in this pathway was significantly up-regulated (Supplementary Table S5 and Supplementary Figure S5). These down-regulated genes included both the large and small subunit ribosome proteins. The down-regulated DEGs also included a group of 98 genes with transmembrane domain (Supplementary Table S6), but with only 1.5-fold enrichment. These genes included many involved in nutrient transport across the membrane, as well as several kinases and other genes. The potential molecular pathway(s) PiRV-2 employed to act on *P. infestans* were further explored in the “Discussion” section.

### Distribution of PiRV-2 in *P. infestans*

Since PiRV-2 stimulated sporulation in *P. infestans* and enhanced its virulence, we suspected that it could play a role in late blight epidemics and set out to examine its presence in *P. infestans* population. In a previous study, we screened 22 isolates of *P. infestans* using dsRNA extraction [16]. In the current study, we screened 54 isolates for PiRV-2 using RT-PCR (Fig 4 and not shown). The combined results are detailed in supplementary Table S1 and summarized in Table 1. In North America, the virus was detected in 11 out of 13 isolates in the US-8 lineage and 3 out of 4 isolates in US-22 lineages, but not in 6 isolates belonging to US-11, US-17, US-23 or isolates whose lineages were not determined. From isolates collected outside of the USA, PiRV-2 was not detected in 9 isolates from Estonia, The Netherlands and South America. However, PiRV-2 was detected in 1 out of 41 isolates from Mexico.

**Fig 4.**
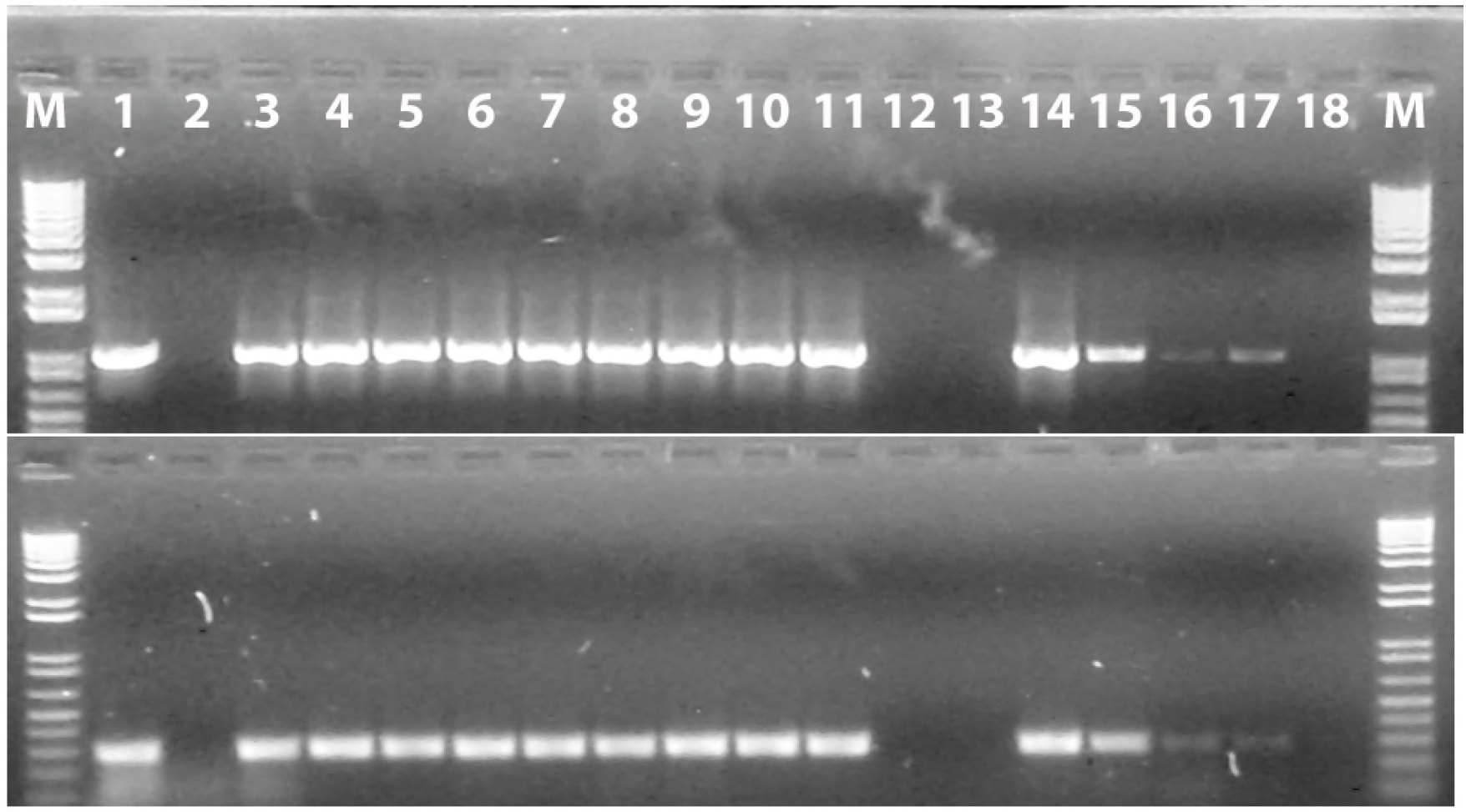
RT-PCR detection of PiRV-2 in *P. infestans* collection. Upper panel, primer pair P24/P27, lower panel, P16/P18. Lane designation: M, 1kb DNA ladder; 1, US940480; 2, negative control; 3, US040009; 4, P8449; 5, P9017; 6, P9051; 7, P9200; 8, P9212; 9, P9219; 10, P10107; 11, P10129; 12, US100001; 13, US100002; 14, US110001; 15, P17707; 16, P17777; 17, P17783; and 18, P17785.

**Table 1.** PiRV-2 distribution in *Phytophthora infestans*.

| Location/lineage |  | # of isolates | PiRV-2 positive | PiRV-2 negative |
| --- | --- | --- | --- | --- |
| USA | US-8 | 13 | 11 | 2 |
|  | US-17 | 1 | 0 | 1 |
|  | US-11 | 1 | 0 | 1 |
|  | US-22 | 4 | 3 | 1 |
|  | US-23 | 1 | 0 | 1 |
|  | ND | 3 | 0 | 3 |
| South Africa |  | 1 | 0 | 1 |
| The Netherlands |  | 5 | 0 | 5 |
| Estonia |  | 3 | 0 | 3 |
| Mexico |  | 41 | 1 | 40 |

### PiRV-2 is faithfully transmitted through asexual reproduction

To determine PiRV-2 transmission through asexual reproduction, single sporangia cultures were obtained from isolate US940480. RT-PCR was used to assay PIRV-2 in 29 single-sporangia cultures, and the virus was detected in all of them (Fig. 5).

**Fig 5.**
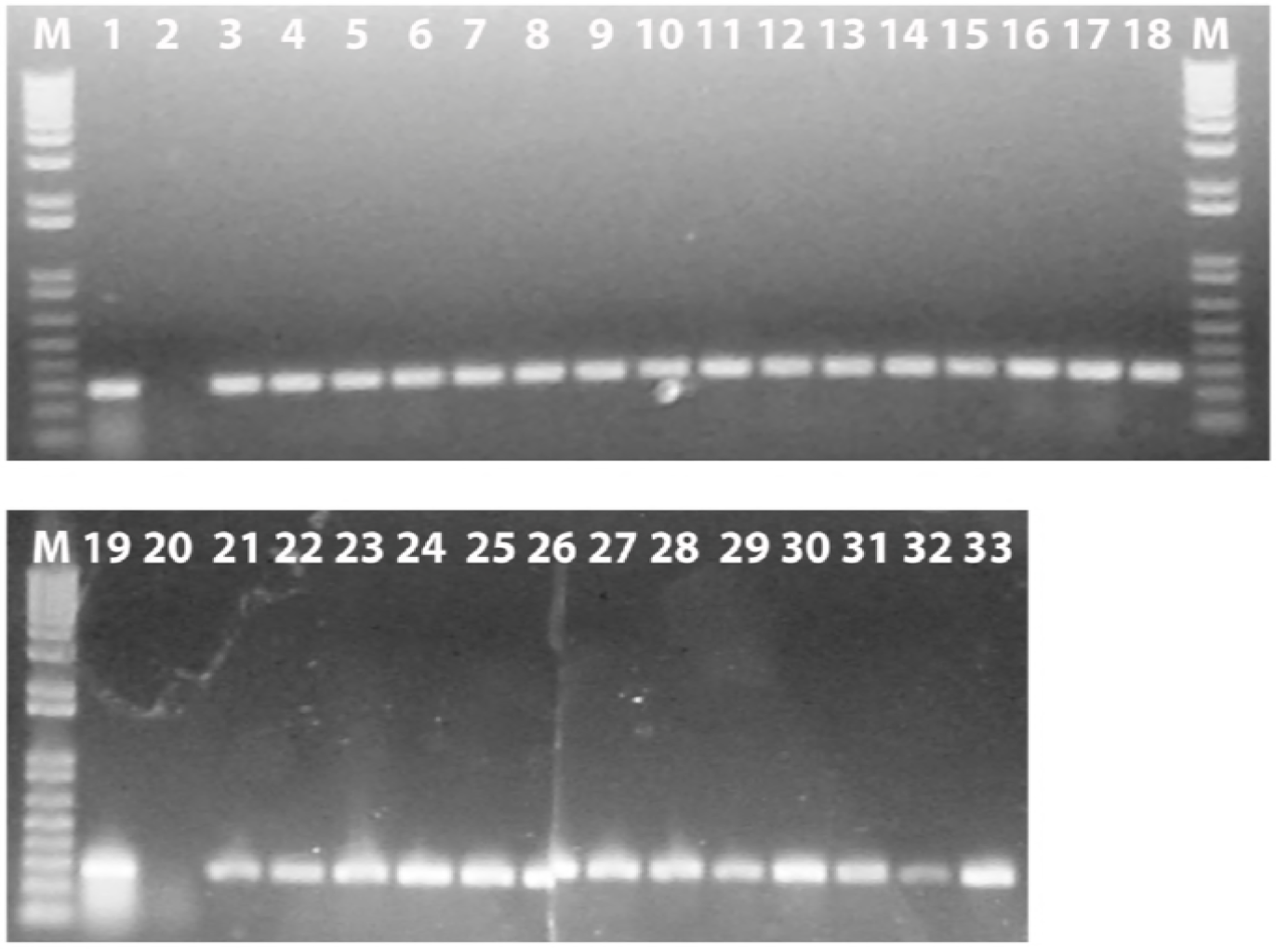
PiRV-2 transmission through sporangia. M, 1 kb DNA size marker, 1 and 19, US940480; 2 and 20, negative controls; and 3<18 and 21<33, single sporangia cultures of US940480.

## Discussion

In this study, we characterized PiRV-2, a novel RNA virus, in the potato and tomato late blight pathogen, *P. infestans*. Efforts to purify VLPs were not successful, but this was not surprising. It is not uncommon for viruses infecting the lower eukaryotes such as oomycetes and fungi to lack protein capsids since most do not infect through an extracellular route. Whether the PiRV-2 dsRNA constitutes the “virus genome” or the replicative form of a single-stranded RNA virus is a matter of definition: viruses in the fungus-infecting family *Hypoviridae* have no capsid protein, but their replicative form dsRNA is encapsulated with an RdRp complex in lipid vesicles [29]. Genomes of these viruses are organizationally and phylogenetically most similar to positive-sense RNA viruses; therefore some virologists consider them to be ssRNA viruses while others consider them to be dsRNA viruses because of its encapsulation in vesicles. Although the RdRp region of PiRV-2 showed some similarity to RdRp_1 domain, which suggests a single-strand RNA genome, the similarity was only marginal. In fact, the RdRp in PiRV-2 was only identified by looking for the three well-conserved motifs in the palm domain of RdRp. Both the 5’ UTR (GUUAAA, 83% AU) and the 3’ UTR (AUUAAGAUGCAUAAUUAAACACAACUUAAGC, 74% AU) of PiRV-2 were AU-rich while the complete sequence had a 52% AU content. An AU-rich 5’ UTR on the plus strand is a general feature of dsRNA viruses and is the region where the dsRNA separates during virus replication as the polymerase enters [30]. No other similarity was observed between PiRV-2 and previously reported viruses. Phylogenic analysis confirmed that PiRV-2 does not belong to any known virus families or groups. We conclude that PiRV-2 belongs to a new virus family yet to be described.

Many viruses of lower eukaryotes are asymptomatic or cause only mild symptoms in their hosts, but others cause observable symptoms [19, 31, 32]. In phytopathogenic fungi, a small number of viruses can attenuate the virulence of their hosts, resulting in a phenomenon called hypovirulence. The best example of hypovirulence was illustrated by Cryphonectria hypovirus 1 [33]. L1 dsRNA in *Nectria radicicola*, the causal agent of ginseng root rot, showed the opposite effect. This virus stimulated sporulation and increased virulence of *N. radicicola* [34]. Alternaria alternata chrysovirus 1 (AaCV1) showed two-sided effects. AaCV1 was isolated from *Alternaria alternate* Japanese pear pathotype, which infects the susceptible pear cultivar ‘Nijisseiki’, produces AK-toxin and causes Alternaria black spot on the leaves. On the one hand, high titer of AaCV1 severely impaired the growth of *A. alternata;* on the other hand, it increased AK-toxin production and enhanced virulence of this fungus [35].

In this study, PiRV-2 appears to have shown similar effects to its host as L1 dsRNA in *N. radicicola*, in that it stimulates sporangium production and enhances virulence. The sporangium plays an important role in the late blight disease cycle [36]. Sporangia are dispersed via air or water to healthy tissue to initiate new infections. Within a few days of inoculation, the pathogen produces a macroscopically visible lesion from which many sporangia can be produced and each is capable of initiating another cycle of pathogenesis.

Given the effects of PiRV-2 on *P. infestans*, it’s reasonable to suspect that this virus could play a role in late blight epidemics. Survey of *P. infestans* collection showed that this virus was present in most isolates of the US-8 clonal lineage of this pathogen. The US-8 lineage was first detected in the United States in 1992 and soon spread to most potato production areas in North America [13]. From the mid-1990s until 2009, US-8 had been the dominant lineage in the USA. Recently (2017), US-23 has been dominant, but US-8 still constituted a substantial portion of *P. infestans* isolated from potato in 2016 and 2017 [4]. The virulence of *P. infestans* depends on many factors, such as inoculum production, temperature adaptation and host preference [15]. We suggest that PiRV-2 may have been a contributing factor to the success of the US-8 lineage. PiRV-2 was found in one of 46 isolates from Mexico. It is assumed that the US-7 and US-8 lineages migrated to the United States in 1992 through the trade of infected plant material [37]. Likely one of the strains carried by the infected material harbored PiRV-2 and led to the dominance of US-8 for several decades in many potato fields in North America.

Clonal lineage US-22 was detected in 2007 and US-23 and US-24 were reported in 2009 [14]. The late blight epidemics in 2009 was mostly caused by US-22. Infected tomato transplants infected by US-22 were distributed by a single supplier and sold by major distributors to many households. It caused wide-spread infection in garden tomatoes, which then spread to commercial tomato and potato fields [12]. Three out of four US-22 isolates examined in this study contained PiRV-2. Further population and epidemiology studies are needed to understand the role of PiRV-2 in late blight epidemics.

At the transcriptome level, the strain harboring PiRV-2 had significantly lower expression of many ribosome protein genes comparing to its isogenic, virus-free counterpart. The ribosome is the core of protein translation machinery. In *Escherichia coli, Neurospora crassa* and *Saccharomyces cerevisiae*, the expression of ribosome protein genes is proportional to growth rate and down-regulated by nutrient limitations [38-40]. In the diploid yeast *S. cerevisiae*, nitrogen starvation induces sporulation and down-regulation of more than 40 ribosome protein genes. When nitrogen is replenished, these genes return to normal expression level and the yeast resumes vegetative growth [41, 42]. While the sporulation in *S. cerevisiae* is a sexual process, resulting in haploid progenies from diploid parent, nitrogen starvation also induces asexual conidiation in fungi *N. crassa* and *Aspergillus nidulans* [43, 44]. There are five annotated ammonium transporter genes in *P. infestans*. Among these, the three genes with the highest expression level were significantly down-regulated by PiRV-2 (Supplementary Table S7). In yeast, down-regulation of ribosome protein genes can also be induced through amino acid deprivation [45]. Among the 34 amino acid permease genes in *P. infestans*, eight were significantly down-regulated by PiRV-2 and none was up-regulated (Supplementary Table S7).

The main function of histone proteins in eukaryotic cells is to package and order DNA. When *S. cerevisiae* is induced into sporulation through nitrogen starvation, the amount of histone proteins per cell increases many folds in the early hours, followed by increase of DNA synthesis [46]. The four major families of histones, H2A, H2B, H3 and H4, form the nucleosome core on which DNA wraps around, while H1 links individual nucleosome together to form higher order structure. Multiple histone protein genes were highly induced by PiRV-2, including all five major histone families (Supplementary Figure S4A and Supplementary Table S4).

In our study, none of the DNA polymerase genes was significantly affected by PiRV-2 but DNA content was not quantified. Genes in one of the up-regulated groups, the seven genes with EGF-like domain, have functions predicted to affect DNA synthesis (Supplementary Figure S4B). EGF-like domains are usually found in the extracellular portion of membrane proteins or in proteins that are secreted. Five of the seven up-regulated genes were predicted to encode extracellular proteins. In animal cells, binding of an EGF-like domain to a cell surface receptor is essential for activation of tyrosine kinase in the receptor cytoplasmic domain, which initiates a signal transduction and results in DNA synthesis and cell proliferation [47, 48].

Previous studies have identified two genes with regulatory functions that are critical for sporangium production in *P. infestans*. A G protein β-subunit (*Pigpb1*) is induced in nutrient starved medium prior to the onset of sporangium formation, and silencing of this gene results in drastic decrease in sporangia production and dense growth of aerial mycelium [49]. *Picdc14* is a cell-cycle phosphatase and expressed only in sporangiophore initials, sporangiophore and sporangia[50]. Silencing of *Picdc14* also impairs sporangium production. *Picdc14* is also induced by nutrient starvation and it is assumed to act after *Pigpb1* because silencing of the latter down-regulates its expression [49, 51]. In our study, *Pigpb1* (PITG_06376) was not significantly affected by PiRV-2. The expression of *Picdc4* (PITG_18578) was up-regulated by 4-fold by PiRV-2 but there was high variability among the replicates (LFC = 1.97, *P* = 0.048, padj = 0.215).

The analysis above suggests a potential molecular mechanism for stimulation of sporulation in *P. infestans* by PiRV-2 (Fig. 6). The virus, directly or indirectly, down-regulates the expression of ammonium and amino acid transporter genes, resulting in nitrogen and amino acid deprivation. This leads to decrease in many ribosome proteins and increase of histone proteins, primes *P. infestans* for DNA synthesis through up-regulation of genes with an EGF-like domain, and possibly up-regulates the expression of *Picdc14*. These molecular processes result in less vegetative growth in strains containing PiRV-2 (sparse aerial mycelium) and more cell proliferation (increased sporangium production) in those strains.

**Fig 6.**
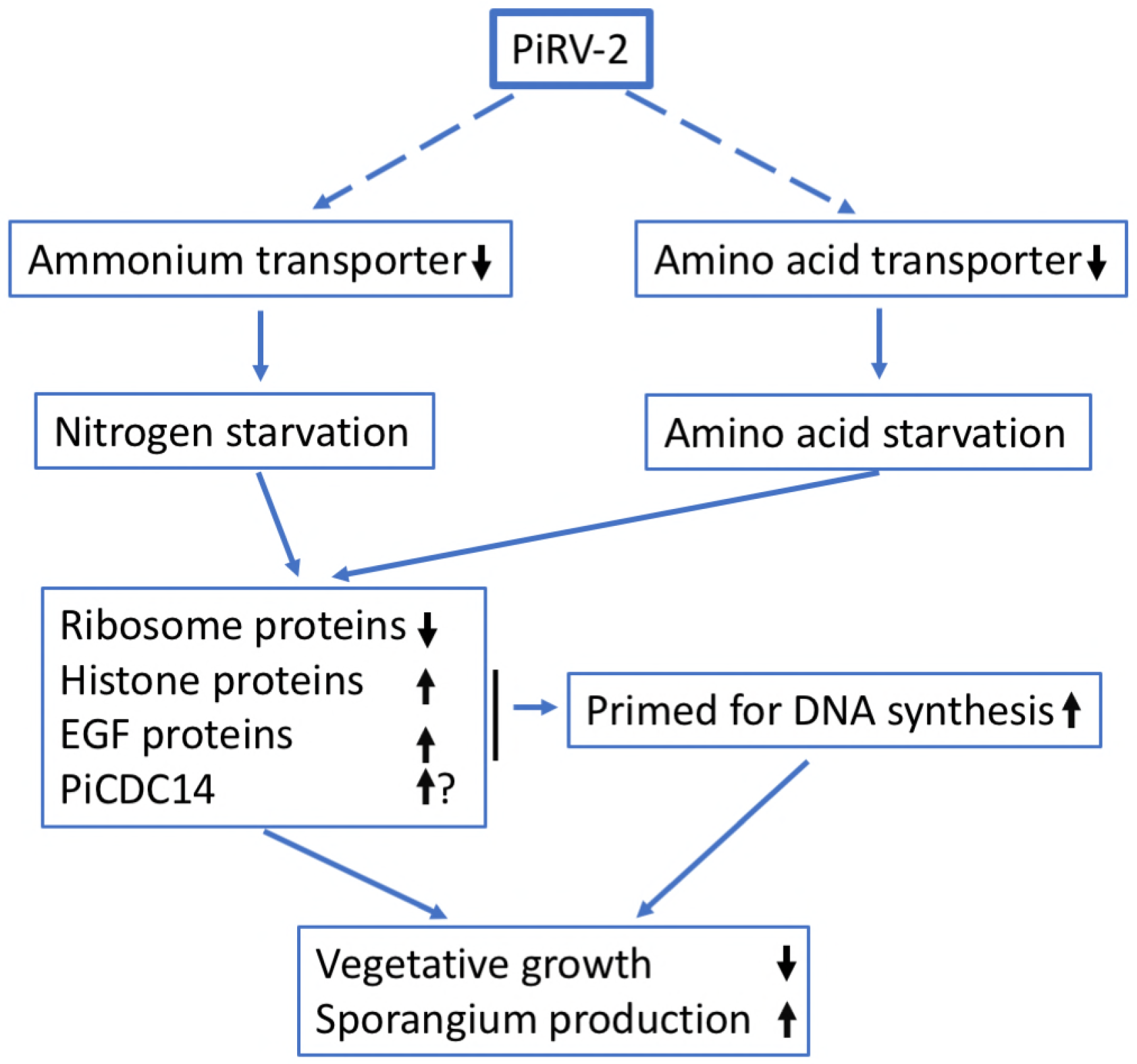
Potential molecular mechanism for PiRV-2-associated stimulation of sporangium production in *P. infestans*.

## Materials and Methods

### Cultures of *P. infestans*

*P. infestans* isolates used in this study and their relevant information are listed in Supplementary Table S1. The isolates were maintained on rye agar. For mycelium production, the isolates were grown in pea broth at 18 °C in darkness.

### RNA extraction

DsRNA was extracted from approximately two-week old mycelium with CF-11 cellulose following the method of Morris and Dodds [52] modified by Tooley et al [53]. DNaseI and S1 nuclease were used as previously described [16] to remove traces of genomic DNA and ssRNA, respectively. Total RNA was extracted from mycelium using RNeasy Plant Mini Kit (Qiagen) according to manufacturer’s instructions.

### Primers

Primers and adapters used throughout this study are listed and described in Supplementary Table S2.

### cDNA cloning, sequencing, and sequence analysis

PiRV-2 was previously found in two isolates of *P. infestans*, US940480 and US040009. The dsRNA from isolate US940480 was used as template for PiRV-2 sequencing. A random cDNA library was constructed as previously described [16], and their sequences were assembled into contigs. Sequence-specific primers were then used in various combinations in reverse transcription (RT)-PCR to confirm sequences of these contigs and to link the contigs in correct order. Three independent rounds of RT-PCRs were performed The primers were designed based on sequences of random clones and sequences of clones from previous round(s) of RT-PCR.

To obtain terminal sequences, a 5’-phosphorylated and 3’-blocked adapter (5’ PO_4_- AGGTCTCGTAGACCGTGCACC-NH2-3’) was ligated to the 3’ end of each strand of the dsRNA using T4 RNA ligase (Promega) [54, 55]. Primer Padapter (5’-GGTGCACGGTCTACGAGACCT-3’), reverse complement to the adapter, was used in combination with sequence-specific primers to amplify terminal regions (Supplementary Figure S1). Superscript III reverse transcriptase and Platinum *Taq* high fidelity enzyme mix (Invitrogen) were used for RT-PCR following manufacturer’s instructions. Amplified fragments were cloned into pCR2.1-TOPO vector (Invitrogen). One or more clones from each RT-PCR were sequenced. All sequencing was conducted using Big Dye Terminator chemistry. Sequences were assembled using Sequencher (Gene Codes Corp.). After PiRV-2/US940480 was sequenced, PiRV-2/US040009 was amplified into two overlapping RT-PCR fragments, and they were purified and sequenced directly (Supplementary Figure S1). The virus genome sequences were deposited in GenBank (accession #s: MH013270 and MH013271).

ORF Finder (http://www.ncbi.nlm.nih.gov/gorf/gorf.html) was used to identify potential ORFs using standard genetic codes. The nucleic acid and deduced protein sequences of PiRV-2 were used to search various databases as described in the “Results” section using default settings. Identification of potential signal peptide and transmembrane regions were carried out using SignalP 4.1 server [56] and TMHMM server v.2.0 [57], respectively. Sequence alignment of RdRps was conducted using the Muscle program in MEGA v.6.0 [58] with visual inspection, and a neighbor-joining tree was constructed and bootstrapped with 1000 replicates.

### PiRV-2 curing

Isolate US940480 was grown on a 9-cm diameter rye agar plate containing 50 mg/ml Ribavirin (SigmaAldrich) at18 °C in the dark. When the colony diameter was approximately two-thirds of the diameter of the plate, a hyphal tip was taken from the edge with the aid of a dissecting microscope and transferred to a new plate. This process was repeated and PiRV-2 curing in the hyphae-tipping cultures was periodically examined using dsRNA extraction and RT-PCR. For this purpose, the hyphae-tipping cultures were grown in pea broth without Ribavirin to produce mycelium. DsRNA and total RNA were extracted from the mycelium as described above. dsRNA samples were run on 1% agarose gel, stained with ethidium bromide and photographed under uv light. SuperScript III one-step RT-PCR system (Invitrogen) was used in RT-PCR, with total RNA as template and primer pairs P05/P07 or P24/P27 (supplementary Table S2) in separate reactions and 40 amplification cycles. The RT-PCR products were separated on agarose gels, stained and photographed as described above. The hyphal-tipping cultures derived from US940480 in which PiRV-2 was cured were named US940480/PiRV-2C1 and US940480/PiRV-2C2.

PiRV-2 transmission through anastomosis. Agar plug from the edge of actively growing US940480 was placed in close proximity with an agar plug from US940480/PiRV-2C1 or US940480/PiRV-2C2 on rye agar plates and grown at 18 °C in the dark. After two weeks, an agar plug was taken from the side of PiRV-2-cured cultures and placed on a fresh rye agar plate. RT-PCR was used to detect PiRV-2. The resultant cultures from US940480/PiRV-2C1 and US940480/PiRV-2C2 into which PiRV-2 was re-introduced were named US940480/PiRV-2T1 and US940480/PiRV-2T2, respectively.

### Phenotype of PiRV-2

Agar plugs from US940480, US940480/PiRV-2C1, US940480/PiRV-2C2, US940480/PiRV-2T1 and US940480/PiRV-2T2 were taken from the edges of actively growing cultures with a #5 cork borer and transferred to the center of fresh 9-cm diameter rye agar plates. The plates were incubated at 18 °C in the dark. Colony diameter was measured at day 5, 7 and 9 after inoculation by averaging two measurements at 90 degree difference. Two weeks after incubation, when the plates were fully covered, they were photographed. Sporangia were harvested by flooding the colonies with 10 ml distilled water and rubbing with a sterile glass rod, and concentrated to 2 ml by low-speed centrifugation. Sporangia concentration was determined using a hemocytometer. The experiment was conducted twice with three plates per isolate in each experiment.

### Pathogenicity test

Sporangia were harvested from 2-week old cultures of US940480 and US940480/PiRV-2C1 as described above. Sporangia concentration was adjusted to 3.5×10^5^ per ml with the aid of a hemocytometer. Leaves were excised from greenhouse-grown Red La Soda potato plants near flowering stage. A 50 ul aliquot of sporangia suspension (17,500 sporangia) was placed in the center of individual leaves. Control leaves were inoculated with sterile H_2_O. Three leaves were used per treatment. The inoculated leaves were placed in a moisture chamber and incubated in a growth chamber at 16 °C and 12-hr light. Lesion diameter was measured by averaging two measurements at 90 degree difference after 7 days.

### Transcriptome sequencing and analysis

Agar plugs from the edge of actively growing cultures of isolates US940480 and US940480/PiRV-2C1 were transferred to rye agar plates overlaid with cellophane, with three plates per isolate. After incubation at 18 °C in dark for 1 week, mycelium from each plate was harvested and total RNA was extracted using RNeasy Plant Mini Kit (Qiagen). An Illumina TruSeq mRNA library was made for each sample according to manufacturer’s instructions. The six libraries, each with distinct barcode, were pooled and sequenced on Illumina Miseq platform for two runs, with the first run at 1x 65 bp and the second run at 1x 66 bp.

Read quality was examined using FastQC [59]. The adapter sequences were trimmed using fastx_clipper program in the FASTX-Toolkit [60]. Reads less than 25 bp after trimming were removed from downstream analysis. The sequence reads were first mapped to PiRV-2 using TopHat v2.1.0 [61] and Bowtie2 v2.3.3.1 [62] to determine if the libraries contained PiRV-2-derived sequences. The genome assembly and annotation of *P. infestans* strain T30-4 [28] were downloaded from GenBank (assembly ASM14294v1). The sequence reads were then mapped to the genome of *P. infestans* and the read counts that mapped to individual protein-coding genes were obtained using HTSeq-count [63]. Differentially expressed genes were analyzed using DESeq2 [64]. Functional analysis of differentially expressed genes were conducted using DAVID version 6.8 [65] with default settings. The transcriptome data were submitted to GenBank (BioProject accession #: PRJNA437643).

### Detection of PiRV-2 in *P. infestans* population

A panel of 54 isolates was maintained on rye agar. They were transferred to pea broth for mycelium growth. Total RNA was extracted from mycelium. Two RT-PCR reactions, with primer pairs p16/p18 and p24/p27, respectively, were used to detect PiRV-2 in each isolate.

### PiRV-2 transmission through sporangia

Sporangia were harvested from 2-week old cultures of US940480 as described above. The concentration of the sporangia suspension was adjusted to ~1,000 sporangia/ml with the aid of a hemocytometer. A rye agar plate was inoculated with 0.3 ml sporangia suspension, spread with a sterile glass rod, air-dried under a hood, and incubated at 18 °C in the dark. Germinating single sporangia were transferred to fresh rye agar plates with the aid of a dissecting microscope to obtain single-sporangium cultures of US940480. To determine the presence of PiRV-2, single-sporangia cultures were grown in pea broth and total RNA was extracted from mycelium. RT-PCR with primer pair p16/p18 was used to detect PiRV-2.

Mention of trade names or commercial products in this publication is solely for the purpose of providing specific information and does not imply recommendation or endorsement by the U.S. Department of Agriculture. USDA is an equal opportunity provider and employer.

## Supporting information

### Supplementary Figures

**Table S1.** Isolates of *Phytophthora infestans* used in the current study

**Table S2.** Anotation of primers and adapters used in the current study

**Table S3.** Impact of PiRV-2 on host gene expression

**Table S4.** Effect of PiRV-2 on expression of histone protein genes and histone modifying enzyme genes

**Table S5.** Effect of PiRV-2 on expression of genes in KEGG ribosome pathway

**Table S6.** Transmembrane proteins down-regulated by PiRV2

**Table S7.** Effect of PiRV-2 on the expression of ammonium transpoter and amino acid permease genes

